# Auxin and tryptophan trigger common responses in the streptophyte alga *Penium margaritaceum*

**DOI:** 10.1101/2024.12.06.627236

**Authors:** Vanessa Polet Carrillo-Carrasco, Martijn van Galen, Jochem Bronkhorst, Sumanth Mutte, Joris Sprakel, Jorge Hernández-García, Dolf Weijers

## Abstract

Auxin is a signaling molecule that regulates multiple processes in the growth and development of land plants. Research gathered from model species, particularly *Arabidopsis thaliana*, has revealed that the nuclear auxin pathway controls many of these processes through transcriptional regulation. Recently, a non-transcriptional pathway based on rapid phosphorylation mediated by kinases has been described, complementing the understanding of the complexity of auxin-regulated processes. Phylogenetic inferences of both pathways indicate that only some of the components are conserved beyond land plants. This raises fundamental questions about the evolutionary origin of auxin responses and whether algal sisters share mechanistic features with land plants. Here we explore auxin responses in the unicellular streptophyte alga *Penium margaritaceum*. By assessing physiological, transcriptomic and cellular responses we found that auxin triggers cell proliferation, gene regulation and acceleration of cytoplasmic streaming. Notably, all these responses are also triggered by the structurally related tryptophan. These results identify shared auxin response features among land plants and algae, and suggest that less chemically specific responses preceded the emergence of auxin-specific regulatory networks in land plants.

## Introduction

Auxin (Indole-3-acetic acid or IAA) is a small signaling molecule that regulates various aspects of plant growth and development including cell division, growth (expansion, elongation) and differentiation [Reviewed in ^1,2^]. Studies in *Arabidopsis* have demonstrated that auxin responses are contex-dependent and vary with the cellular environment. For instance, auxin promotes cell growth in hypocotyls through acidification of the apoplast, which stimulates overall hypocotyl elongation ^3^. In contrast, a similar stimulus leads to root growth inhibition ^4,5^. Exploration of auxin responses across land plants has revealed both conserved and unique responses ^4,6,7^. Despite decades of auxin research, the extent of analogous responses across tissues and species, as well as their evolutionary origin remains unclear, in part due to the complexity of auxin-mediated regulation in land plants.

The most studied signaling module regulating auxin responses is the nuclear auxin pathway (NAP)^2^. This pathway operates through transcriptional regulation and the components of the pathway are phylogenetically and functionally conserved along the land plant phylogeny ^8–11^. In addition to transcriptional responses, auxin triggers rapid non-transcriptional responses such as promoting cytoplasmic streaming, disrupting proton transport, inhibiting root growth or cytoskeleton rearrangements ^12–14^. Recent studies in *Arabidopsis* revealed a distinct non-transcriptional pathway involving auxin-triggered global phosphorylation ^6,15^. This response is independent of the NAP ^5^ and instead relies on a phosphorylation cascade mediated by kinases ^16^.

In streptophyte algae, the closest relatives to land plants, only some NAP components are conserved^8,10^, suggesting that this pathway evolved in land plants ^10,17,18^. While the transcription factors mediating auxin response are conserved between algae and land plants ^19^, the remaining components are missing or distantly related to those operating land plants. Endogenous auxin has been detected in several algal species ^20^ and multiple responses to auxin have been reported in streptophyte algae ^21–25^. These studies reveal contrasting effects between single-celled and filamentous streptophyte algal species. In *Klebsormidium nitens* (Klebsormidiophyceae), high concentrations of IAA inhibit cell division and elongation ^23^. Conversely, much lower auxin concentrations promote cell proliferation in the zygnematophyceaen alga *Micrasterias thomasiana* ^24^. Based on these contrasting responses in phylogenetically very distant algal species, it is unclear if there are common auxin responses in algae, and if these are shared with land plants.

Furthermore, given that the NAP is absent from algae, response pathways other than the NAP must exist in these organisms. Indeed, global auxin-triggered phosphorylation is conserved not only across land plants, but in different groups of streptophyte algae, including *Penium margaritaceum, K. nitens and Chara braunii* ^7,16,26^. Inference of the putative biological functions regulated by this pathway indicated both lineage-specific and conserved functions/targets among the tested species. This showed that auxin-mediated phosphorylation signaling is a deeply conserved mechanism in all streptophytes. However, it remains unknown if this mechanism is connected to physiological responses in streptophyte algae as it is in land plant species.

Here, we have explored auxin responses in the unicellular streptophyte alga *Penium margaritaceum*, evaluating physiological, transcriptomic and cellular responses. We have developed a microfluidic system to follow single algal cells in time-resolved auxin responses. Our results indicate that auxin triggers cell proliferation and shifts in the transcriptome towards gene activation, but also a fast response accelerating cytoplasmic streaming, as occurs in land plants. However, these responses largely overlap with those exerted by tryptophan, a structurally similar molecule, and to a lesser extent with the unrelated benzoic acid. Our results support a common effect of auxin in all streptophytes, but also suggests that the highly specific land plant-specific NAP was preceded by a less chemically specific response system.

## Results

### Auxin and tryptophan promote cell growth and division in *Penium margaritaceum*

To assess whether *P. margaritaceum* (*Penium*) exhibits physiological responses to auxin, we applied varying concentrations of auxin to batch cultures. While various small molecules have auxin activity in land plants, only a few naturally occurring auxins have been identified, and among these, IAA is the dominant one [Reviewed in ^27^]. We therefore used IAA in all our assays and refer to it as “auxin” for simplicity. We monitored growth by measuring optical density in cultures over 10 days (**Fig. 1A**). The growth curves of IAA-treated cultures did not differ significantly from the mock-treated (DMSO) controls. This suggests that IAA does not influence *Penium* growth at the population level. However, optical density measurements in batch cultures represent the average response of the entire population, potentially obscuring cell-to-cell variation and responsiveness.

**Figure 1.**
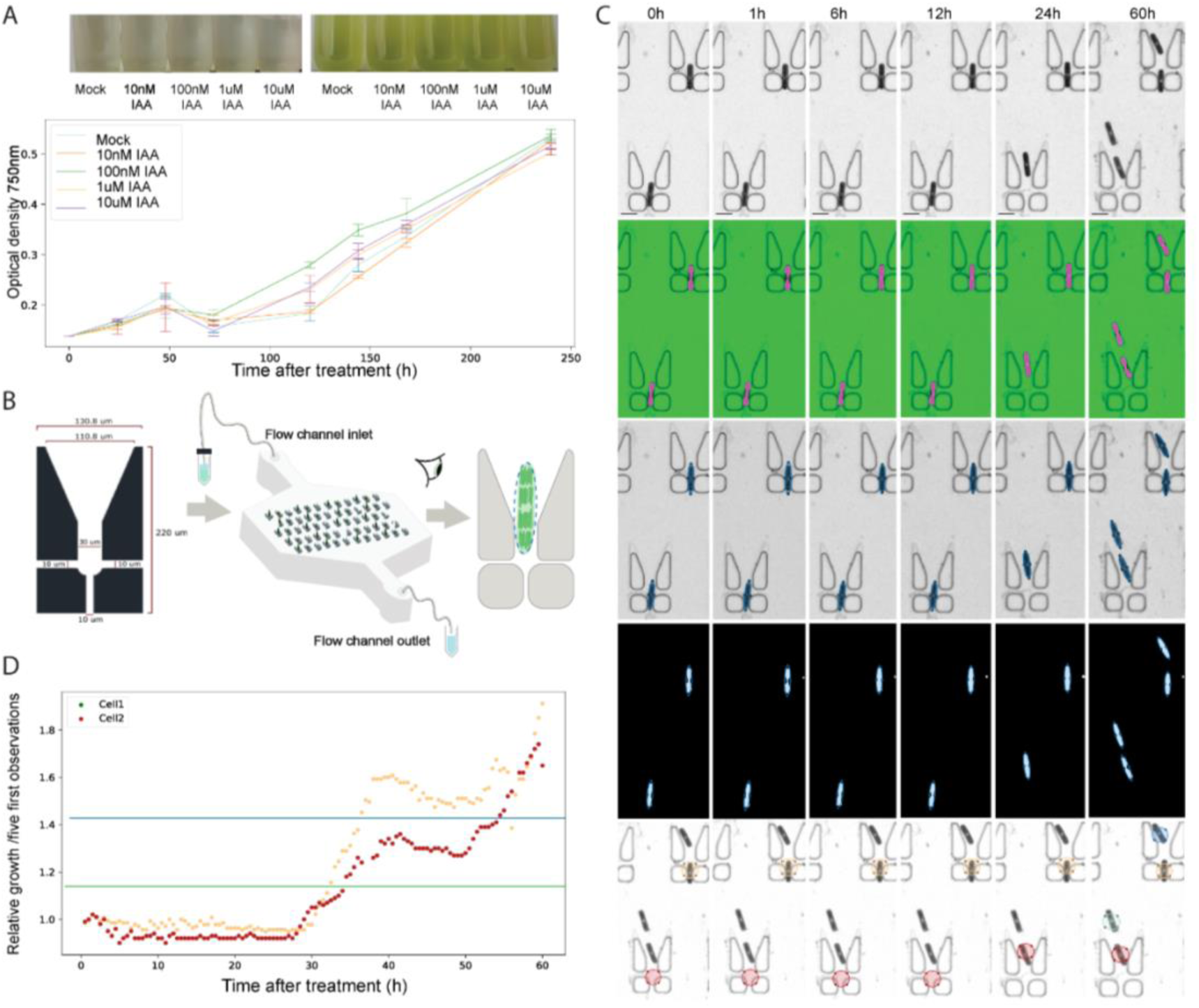
Tracking cellular auxin response in *Penium margaritaceum*. (**A**) Line graphs showing the change in optical density at 750 nm of 50 ml cultures grown for 10 days with different concentrations of IAA. DMSO was used as a mock control. Top row: images of the cultures at 0 and 6 days after inoculation. (**B**) Schematic representation of microfluidic devices used to immobilize *Penium* cells mounted on an inverted microscope for visualization. (**C**) Processing of time series images made using transmitted light microscopy. Images were made in 30-minute intervals over 60 hours. Top to bottom show the raw images, images after binarization, cell segmentation, background removal, and centroid (blue) tracking over time. Each colored circle represents a single cell. Scale bar size 50 µm. (**D**) Traces obtained with the information collected from image processing indicate relative growth curves in periods of 60 hours. Each dot represents a 30-minute interval. The horizontal green/red lines represent the cut-off used to separate growing or dividing cells, respectively.

To investigate the effect of external IAA at the single-cell level, we developed a microfluidics system (see Methods) to trap individual cells. This, coupled with time-lapse microscopy, allowed monitoring growth in real-time (**Fig. 1B**). In addition, we developed a pipeline for automated image analysis, facilitating precise tracking of cell size and division events over time (**Fig. 1C, D**). We used this setup to track cell growth and division over a 60-hour period with media supplemented with IAA, or DMSO as mock for the control group. Cell lengths were measured at 30-minute intervals, and relative growth traces generated based on growth rates between the average length of the initial five observations and subsequent time points (**Fig. 1D**). Individual growth traces were organized by treatment across independent experiments. Under mock conditions, an average of 18% of cells were in an active growth phase per experiment (**Fig. 2A; Fig. S1**), consistent with these cultures not being synchronized, and indicating that less than a quarter of these cells show active growth in our setup (**Video S1**). In contrast, the percentage of actively growing cells in the IAA-treated groups doubled this percentage (41%), showing a physiological effect of the molecule. In line with these results, our analyses of cell division occurrences showed that a third of the IAA-treated cells divide on average, while only 6% of mock-treated cells divide (**Fig. 2B**).

**Figure 2.**
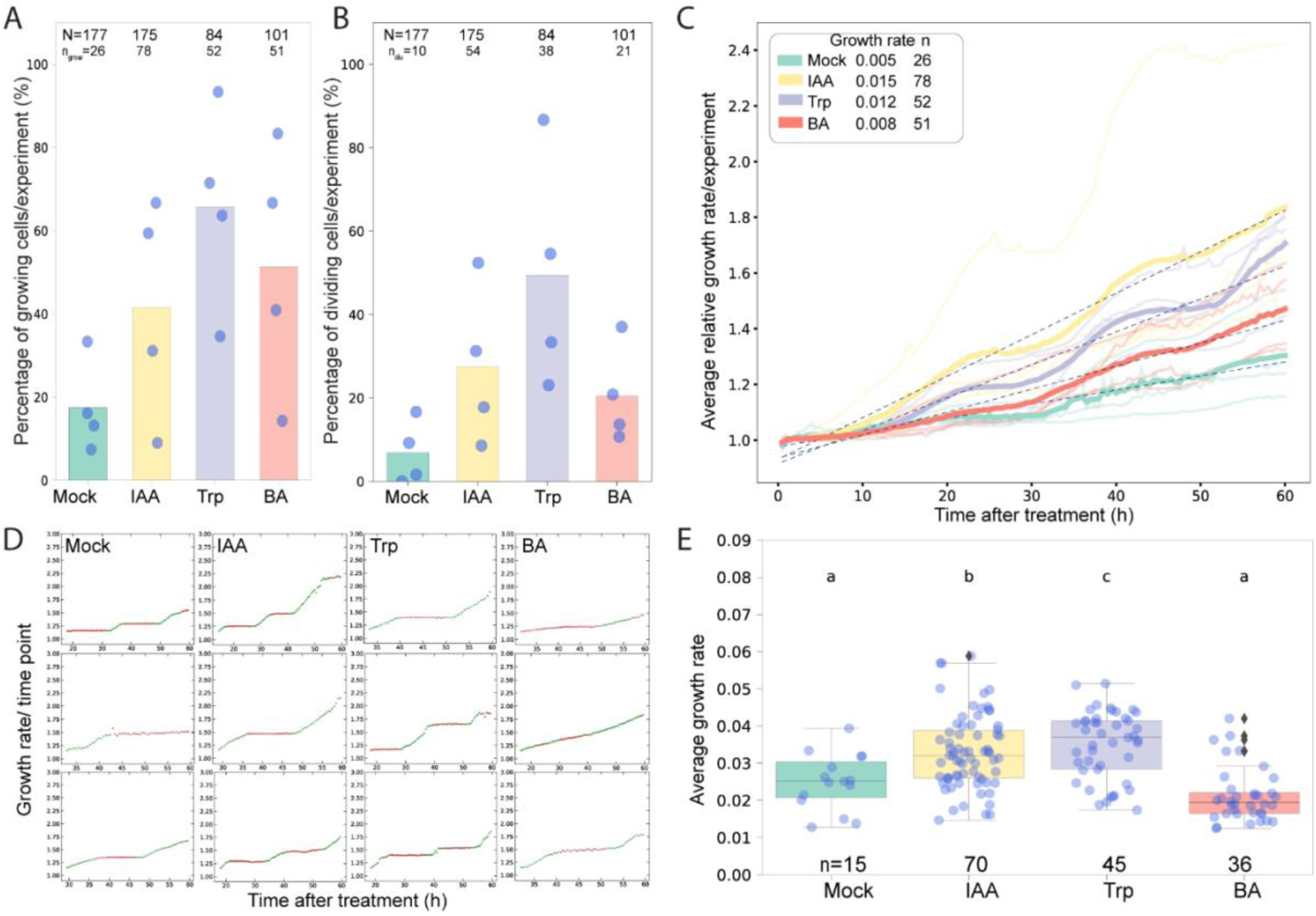
Single-cell responses to auxin, Trp and BA in *Penium*. **(A**) Percentage of growing cells per experiment. Each dot represents the percentage of cells per individual experiment. N, total number of cells evaluated per treatment and n_grow_ number of growing cells. (**B**) Percentage of dividing cells per experiment. n_div_ number of dividing cells. (**C**) Average growth curve of each treatment (thick lines). The trace was constructed using the average growth curve of each independent experiment (thin growth curves). Dashed lines represent the trend line used to estimate the growth rate. (**D**) Single-cell traces adjusted to a trace displacement pipeline. (**E**) Boxplots depict growth rates (µm/h) obtained during growing periods. n= number of cells. Statistical groups were obtained with a one-way ANOVA followed by Tukey’s Post-Hoc test (p<0.05).

We additionally found an effect of IAA on cellular growth rates by following single-cell traces. Cells treated with IAA grew three times faster than those growing under mock conditions (**Fig. 2C; Fig. S1**). Individual traces showed that *Penium* cells undergo a periodic alternation between active growth and growth quiescence. To reduce the possible effect of different treatments interfering or varying these periods and properly compare growth rates, we adapted a recently developed pipeline to measure trace displacement^28^. This way, we could mathematically define such periods, determining only the growth periods to extract growth rates (**Fig. 2D; Fig. S2**). This unambiguously showed that IAA-treated cells also grow faster than those under mock conditions (**Fig. 2E, Table S1**).

IAA in land plants is synthesized from tryptophan in a simple, two-step pathway ^29–31^. However, these enzymes are not consistently present in streptophyte algal genomes ^17,32^. We therefore asked if the response to IAA that we observed in *Penium* is chemically specific to IAA. We treated cells with tryptophan (Trp), as well as with the chemically unrelated organic acid benzoic acid (BA). Both Trp and BA have a pKa similar to that of IAA, and also exist naturally in cells. We found that each of these organic acids had a similar effect in promoting cell growth and division (**Fig. 2A,B**). This was especially true for Trp, even exceeding the effect of IAA on the average number of cells growing (66%) and dividing (45%). The growth rate was also increased upon Trp treatment, but not with BA (**Fig. 2E**). Altogether, these results indicate that *Penium* cells respond to externally applied IAA, but also show that other organic acids can trigger a similar response.

### IAA and tryptophan trigger overlapping transcriptional responses in Penium

Long-term physiological responses such as growth and division are likely mediated by transcriptional reprogramming. To explore the existence of transcriptional responses to IAA in *Penium*, and analyze potential transcriptional overlaps in the responses to IAA, Trp, and BA, we performed whole transcriptome sequencing. Our *P. margaritaceum* strain (NIES-217) data mapped poorly into the available genome annotation of strain Skd8 (<12% reads) ^33^. We, therefore constructed a *de novo* transcriptome assembly for this strain combining short-read and long-read RNA sequencing to which >80% of the reads mapped. Comparison of the assemblies suggests that NIES-217 and Skd8 represent distantly related strains, and potentially even different species (**Fig. S3**). We treated *Penium* cultures with 1µM IAA, Trp or BA for 1 hour, and used the same volume of DMSO as a mock control. We next mapped the reads from each sample to the *de novo* assembly and identified expressed transcripts. Analysis of the global patterns of similarity among treatments and replicates indicate that IAA and Trp induce different transcriptional responses compared to mock. Conversely, BA-treated samples exhibited high variation, with replicates spread across the plot but close to mock samples (**Fig. 3A**).

**Figure 3.**
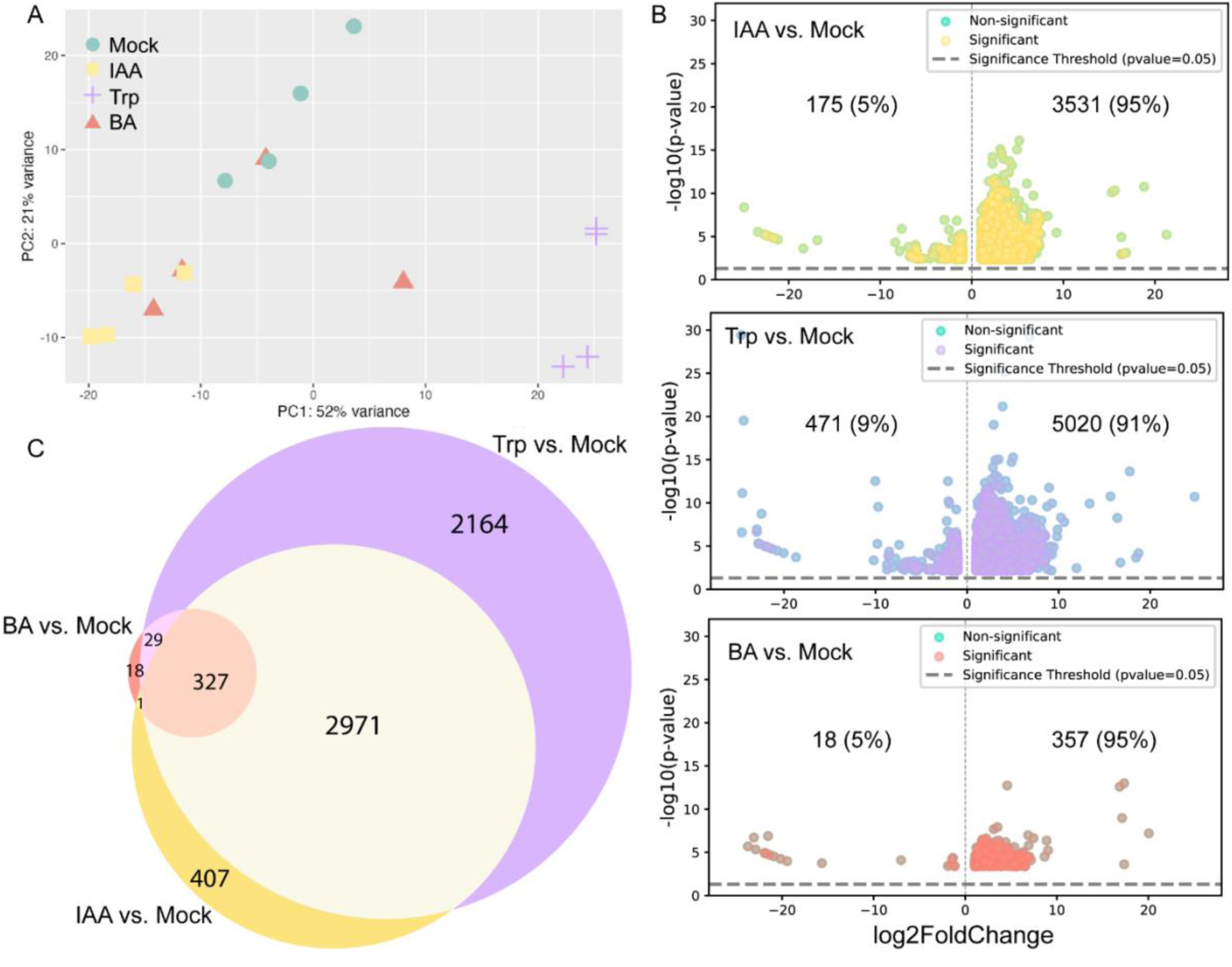
IAA and Trp trigger an overlapping transcriptional response in Penium. **(A**) PCA plot showing the clustering of RNA-seq samples based on gene expression profiles. Axes X and Y indicate the variance in the dataset. (**B**) Volcano plots showing the distribution of differentially expresses genes (DEGs) of cells treated with IAA (yellow), Trp (lilac) or BA (orange) compared with the mock control DMSO. Only statistically significant DEGs are represented (adjusted p-value = 0.05 and log2 fold change=1). (**C**) Venn diagram showing the overlap between the DEGs of each condition. Overlapping regions indicate genes/transcripts shared between conditions.

We identified differentially expressed genes (DEGs) for each treatment compared to the mock. This revealed 3706, 5491, and 375 DEGs in response to IAA, Trp, and BA, respectively (**Fig. 3B**). The majority of DEGs showed upregulation (95%, 91%, and 95%, respectively, log_2_FC >1). When surveying overlap between treatments (**Fig. 3C**), we found a large overlap between IAA and Trp responses (3299 DEGs), representing around 89% of IAA-regulated genes. We observed a small proportion of transcripts that seemed specific to IAA (n=407). However, most IAA-triggered DEGs were also triggered by Trp (n=3299). In addition, we found a substantial set of Trp-specific (n=2164) and a very small number of BA-specific (n=18) DEGs. Thus, this transcriptomic analysis support similar activities of IAA and Trp in *Penium* as observed in growth assays. IAA controls a subset of Trp-regulated genes, reflecting an overlapping response and suggesting that responses are likely not chemically specific to auxin. Gene ontology term enrichment analysis further highlighted the large functional similarity between IAA and Trp treatments (**Fig. S4**). Taken together, these results suggest that algal auxin-regulated responses may represent a broader response to organic acids.

### Auxin and tryptophan promote cytoplasmic streaming

Phospho-proteomic analysis previously showed that IAA can trigger a set of conserved and specific responses within two minutes in land plants and streptophyte algal species, including *Penium* ^16,26,34^. A biological outcome of this rapid response in *Arabidopsis* roots and *Marchantia* rhizoids is an increase in the velocity of cytoplasmic streaming, as observed by motility of mitochondria labelled with Rhodamine 123^16^. To study if this response also occurs in *Penium*, we used the above-mentioned microfluidics system coupled with time-resolved imaging to track the movement of labelled particles. Unfortunately, we found that Rhodamine 123 does not reliably stain motile structures in *Penium*. Instead, we used MitoTracker Orange CMTMRos, which enabled visualization of traceable punctate particles to measure the average speed of particles in cells treated with IAA, Trp, and BA (**Fig. 4A, Video S2**). We observed significant increases in particle motility in cells treated with 1 µM IAA and Trp, while 0.1 µM IAA— the concentration used to evaluate auxin responses in land plants —showed no significant effect (**Fig. 4B**). BA-treated cells exhibited high experimental variability, and no clear effect could be seen.

**Figure 4.**
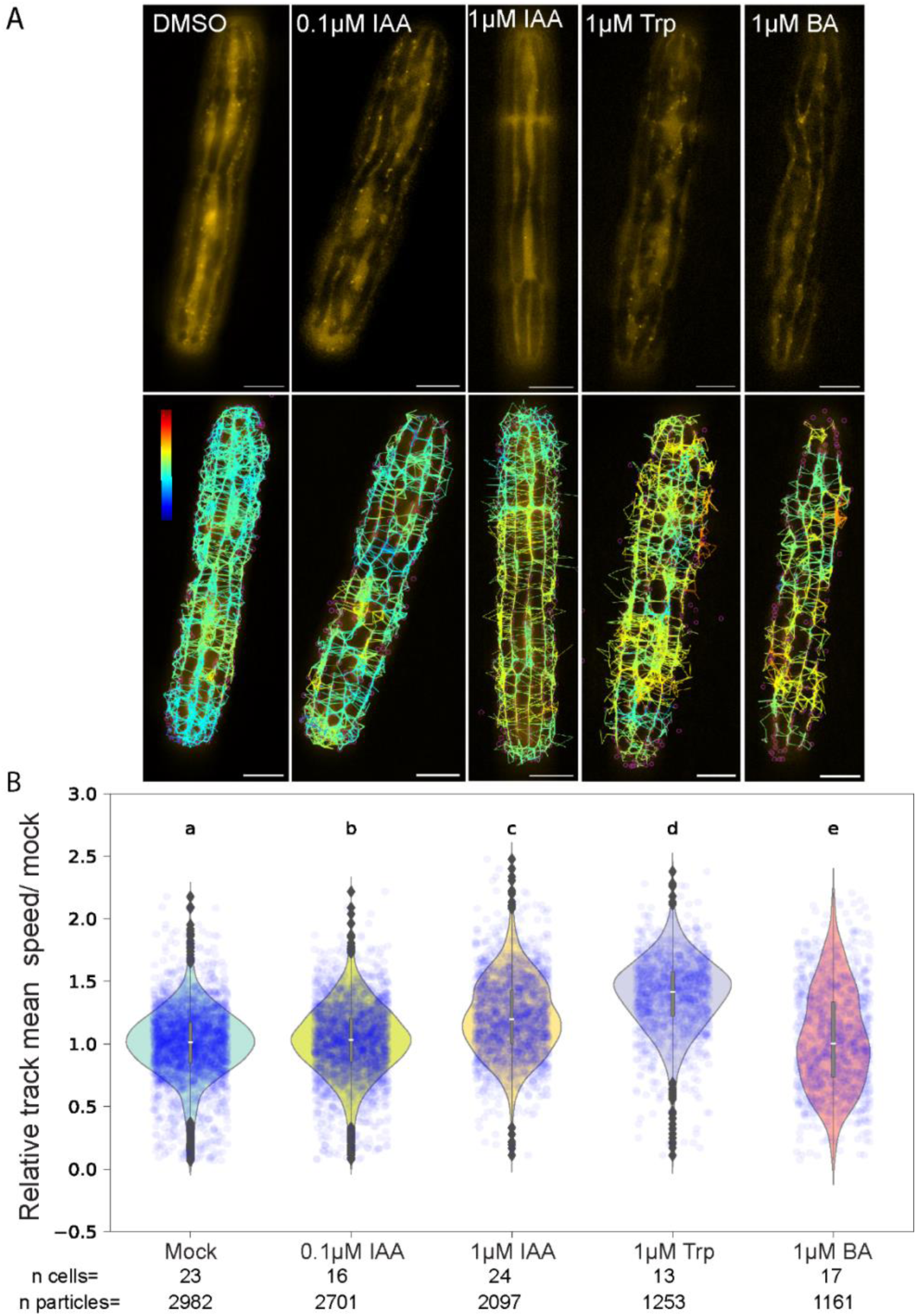
Auxin and tryptophan accelerate cytoplasmic streaming in *Penium.* (**A**) Top: representative single cells labelled with MitoTracker Orange^TM^, immobilized in a microfluidics device and treated for 5 minutes with DMSO, IAA, Trp or BA. Cytoplasmic streaming of particles was recorded over five seconds. Bottom: trajectories obtained from the image analysis. The trajectories were filtered by selecting only particles that can be traced continuously for 3 seconds, colour bar indicates the trackspeed range from 0 to 50 µm/s. (**B**). Violin plots showing the relative average speed of particles in µm/s to the average of the control as a readout of cytoplasmic streaming. Statistical groups were obtained with a one-way ANOVA followed by Tukey’s Post-Hoc test (p<0.05). n cells: total number of cells evaluated per treatment. n particles : total number of tracked particles. Scale bar 10 µm.

These results demonstrate that IAA triggers rapid responses in *Penium*, comparable to the fast auxin-induced responses observed in land plants. However, as with the longer-term responses shown above, this response is elicited by Trp, and not by BA.

## Discussion

In land plants, auxin exhibits context-dependent effects across different tissues and species.^7,9,10,35^. Although responses to auxin have been described in streptophyte algae, no unified response has yet been reported. Using bespoke microfluidic traps and long-term live imaging, we demonstrate that IAA promotes cell growth and division in *Penium*, at least superficially similar to what has been reported in the desmid *M. thomasiana* ^24^. We anticipate that extending the cell-based analysis of responses to IAA across other streptophyte algal species may help define whether there is indeed a common response shared among algae – and between algae and land plants.

The enzymes for canonical IAA synthesis in land plants (TAA/TAR and YUC^29,31,36^) are lacking in many of the streptophyte algal lineages ^18^. However, recent profiling of hormone levels in a range of algal species found that IAA and some of its metabolites can be found to varying degrees ^20^. A key question therefore is what the source of this IAA in algae is. Is there a separate yet dedicated auxin synthesis pathway, is this a catabolic product? While these questions await answers, our work does show that there is a clear response to IAA. We find responses to low concentrations (1 µM) of IAA at cell growth, cell division, transcription, cytoplasmic streaming (this work) and protein phosphorylation ^16^. A key question is, which of these responses are linked within pathways? Is there a single receptor type that relays IAA perception to both rapid and sustained responses, or do the fast and slower responses relay on different systems, as is likely the case in land plants ^6,37^? Given that the NAP appears absent in streptophyte algae^17,18^, it is evident that the mechanism for IAA-triggered transcriptional changes must differ from the one in land plants. Knocking out the single A-class ARF within the Marchantia NAP causes a complete absence of IAA-triggered transcriptional response^38,39^, and thus it is an open question what transcriptional regulators mediate IAA response in *Penium*.

In contrast, several factors required for land plant rapid auxin responses are present in streptophyte algae. These include the ABP1/ABL family of Cupins, as well as TMK1-like receptor-like kinases and RAF-like kinases ^6,16,40^. Thus, the rapid responses in *Penium* – protein phosphorylation and cytoplasmic streaming – may well be part of an ancient, conserved auxin response system. An interesting question is whether the sustained IAA responses also operate through this rapid response system. There are no facile genetic tools for manipulating *Penium* gene function, but through the use of recently developed protein transfection methodology^41^, one could start to functionally dissect the requirements for auxin response in *Penium*.

A key result and surprise was that the responses that IAA induces both on rapid and slow timescales, are equally triggered by tryptophan. Whereas we find some response to Benzoic Acid in the long-term cell growth assays, those are not observed at the level of transcriptome or cytoplasmic streaming. Thus, the responses are chemically specific among organic acids, but IAA responses are in reality not IAA responses, but IAA/Trp responses. This finding has a number of important implications. Clearly, our findings are limited to a single streptophyte algal species, and this may not be representative of the group it belongs to. We therefore cautiously interpret these results, but propose that the chemically highly specific IAA response system in plant plants may have been preceded by or even evolved from, a less specific system that mediated responses to IAA-like molecules. Or perhaps the primary target for this response system is Trp, and IAA is simply a Trp-like metabolite that can use the same response system. The chemical promiscuity of any response system to both IAA and Trp is not unheard of. In fact, precursors of bioactive molecules can act as signals themselves in distant species, as is the case of the ethylene precursor ACC ^42^, the Jasmonic Acid precursor OPDA [Reviewed in ^43^], or the Gibberellic Acid precursor ent-kaurenoic acid ^44^. Furthermore, a recent report indicates that lycophyte jasmonate receptors have a broader range of substrate specificity than euphyllophyte or bryophyte receptors ^45^. One interpretation of the similar responses to IAA and Trp is that Trp is metabolized into IAA, and the responses observed are in fact responses to IAA. The stronger response to Trp and the timeframe do not completely align with this interpretation. Alternatively, the opposite direction – IAA to Trp conversion - is theoretically possible, as bacteria can transform indole into Trp ^46^. Whatever the nature of the response system or synthesis pathway, these results reveal a functional similarity between Trp and IAA, which casts new light onto IAA as a signaling molecule and urges the inclusion of Trp as a control in studies focusing on IAA response.

A key question is what role this IAA/Trp response serves. We distinguish two scenarios. Either IAA and Trp act as metabolites, and the responses are nutritional/metabolic effects. Alternatively, IAA and Trp act as bona fide signals that are perceived by a dedicated receptor, and are relayed to cellular responses. Given the low concentrations used (1 µM), we favor the second interpretation, although the former cannot be excluded. Also, the rapid phosphorylation of a range of signaling proteins, such as protein kinases ^16^ would suggest that this is a signaling response. What, then, are IAA and Trp signals for? This would depend on the source of the molecules. If perception is apoplastic, the source would, by definition, be external in a single-celled organism. In this case, the presence of IAA or Trp in a cell’s vicinity could be (1) a within-species damage signal, (2) an autocrine or paracrine signal (e.g., quorum sensing), or (3) a signal derived from nearby other species. All these options are possible and deserve future investigation.

In summary, our study demonstrates that *Penium* exhibits IAA-mediated cellular, physiological and transcriptional responses, including promoting cell growth, division, and cytoplasmic streaming. The overlapping effects of IAA and Trp suggest potential ancestral roles in environmental sensing and cellular dynamics, distinct from the specialized auxin signaling seen in land plants. These findings lay the groundwork for future studies to unravel the mechanisms of auxin perception and its ecological significance.

## Supporting information

Videao S1

## Resource availability

No new biological materials have been generated for this study. All next-generation sequencing data is available at NCBI under accession numbers PRJNA1192695 and PRJNA1193617. Code is available at https://github.com/PoletC/PhD_thesis.git. Raw data is available from the corresponding authors upon request.

## Acknowledgements

We are grateful to Danilo dos Santos Pereira for help and bioinformatics support, and to lab members for helpful discussions. This work was supported by grants from the Netherlands Organization for Scientific Research (NWO; OCENW.KLEIN.027 to D.W), the European Research Council (ERC; CoG CATCH to J.S), Graduate School VLAG (to M.v.G.), the Human Frontier Science Program (grant RGP0015/2022 to D.W.) and a Marie Skłodowska-Curie Individual Fellowship (MSCA-IF-2020 ref: 101026004 [REOX]) to J.H-G..

## Author contributions

Conceptualization: V.P.C.C. and D.W.; Methodology: V.P.C.C., M.v.G.., J.B., J.S.; Formal analysis: V.P.C.C., M.v.G.., J.B., S.M.; Investigation: V.P.C.C., S.M.; Writing – Original Draft: V.P.C.C.; Writing – Review & Editing: V.P.C.C., J.H.G., D.W.; Visualization: V.P.C.C.; Supervision: J.S., J.H.G., D.W.; Project Administration: V.P.C.C., J.H.G., D.W.; Funding Acquisition: M.v.G.., J.S., J.H.G., D.W..

## Supplemental information

**Figure S1.**
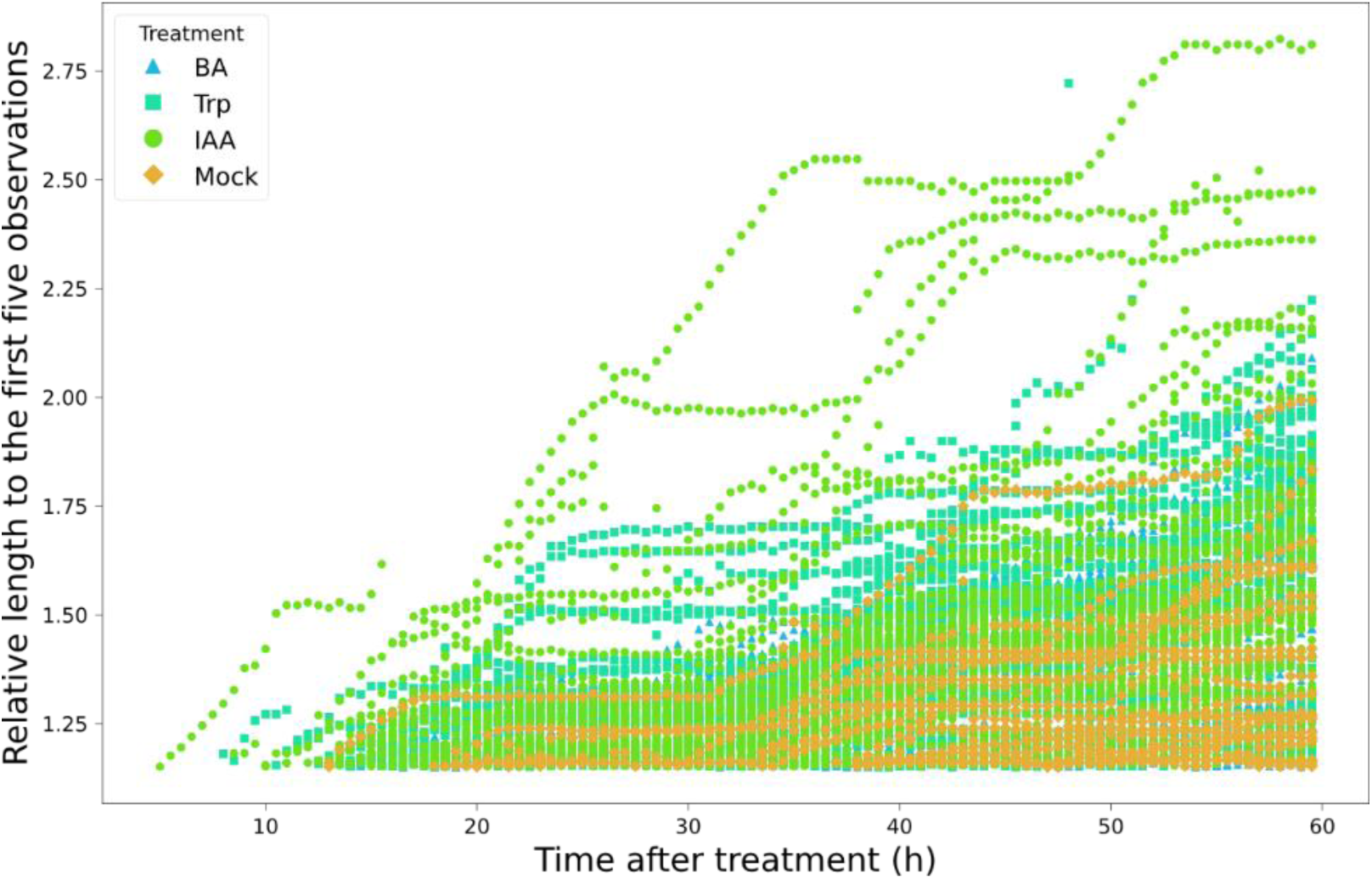
Single-cell growth traces from growth experiments of *Penium* treated with IAA, TRP, BA or DMSO. The triangles represent cell growth traces corresponding to single cells treated with Benzoic acid (BA). Squares represent cell growth traces of cells treated continuously with tryptophan (TRP). Circles represent cell growth traces of cells treated continuously with Indole Acetic Acid (IAA). Diamonds represent cell growth traces of cells treated continuously with DMSO as mock control. Each dot represents a 30-minute interval picture over the course of 60 hours.

**Figure S2.**
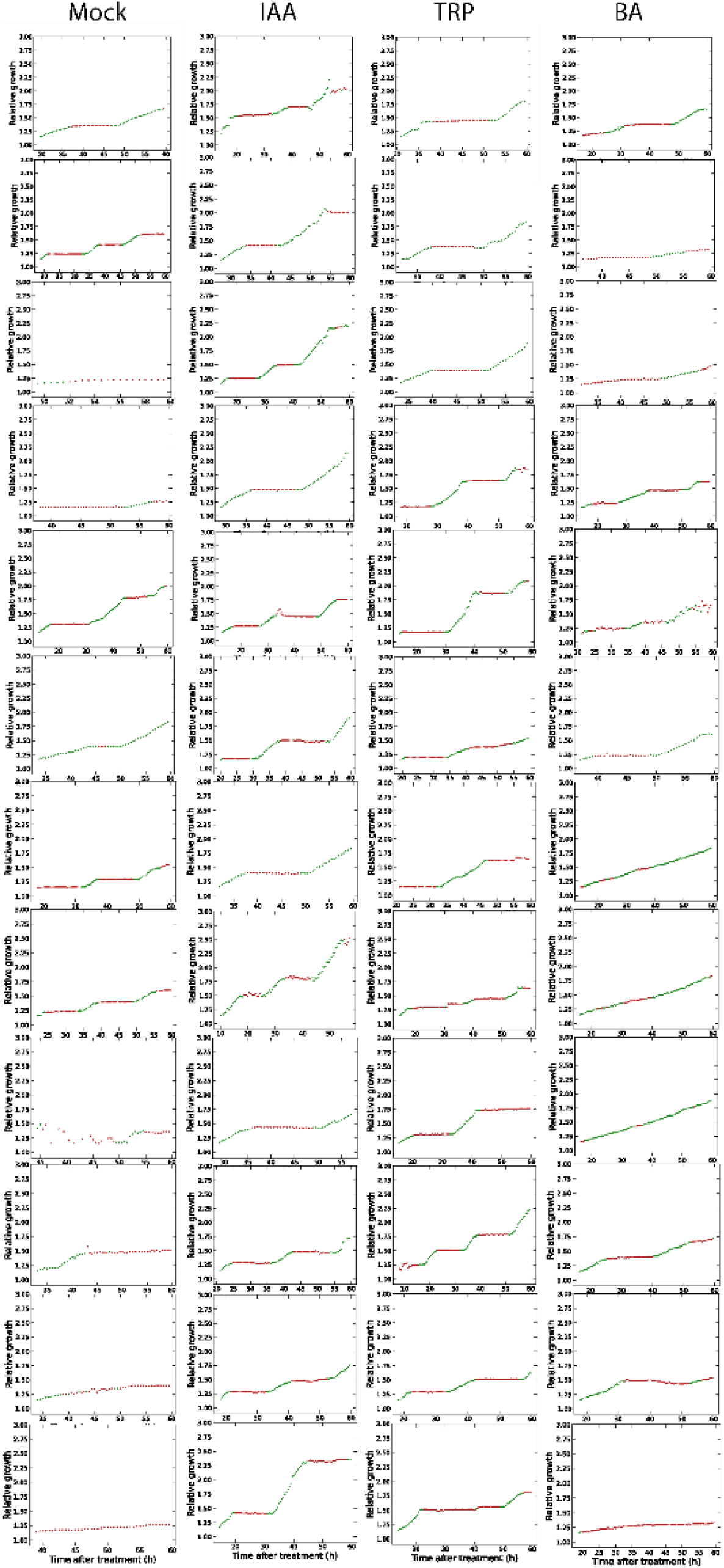
Intermitent growth behaviour of single-cell growth traces. To analyze the intermittent growth behavior of single-cell growth traces, we plotted the relative growth of all evaluated cells and applied a cut-off value to distinguish between growing and non-growing regimes. The chosen cut-off for identifying growing cells was a relative growth rate of 0.012 µm/h.

**Figure S3.**
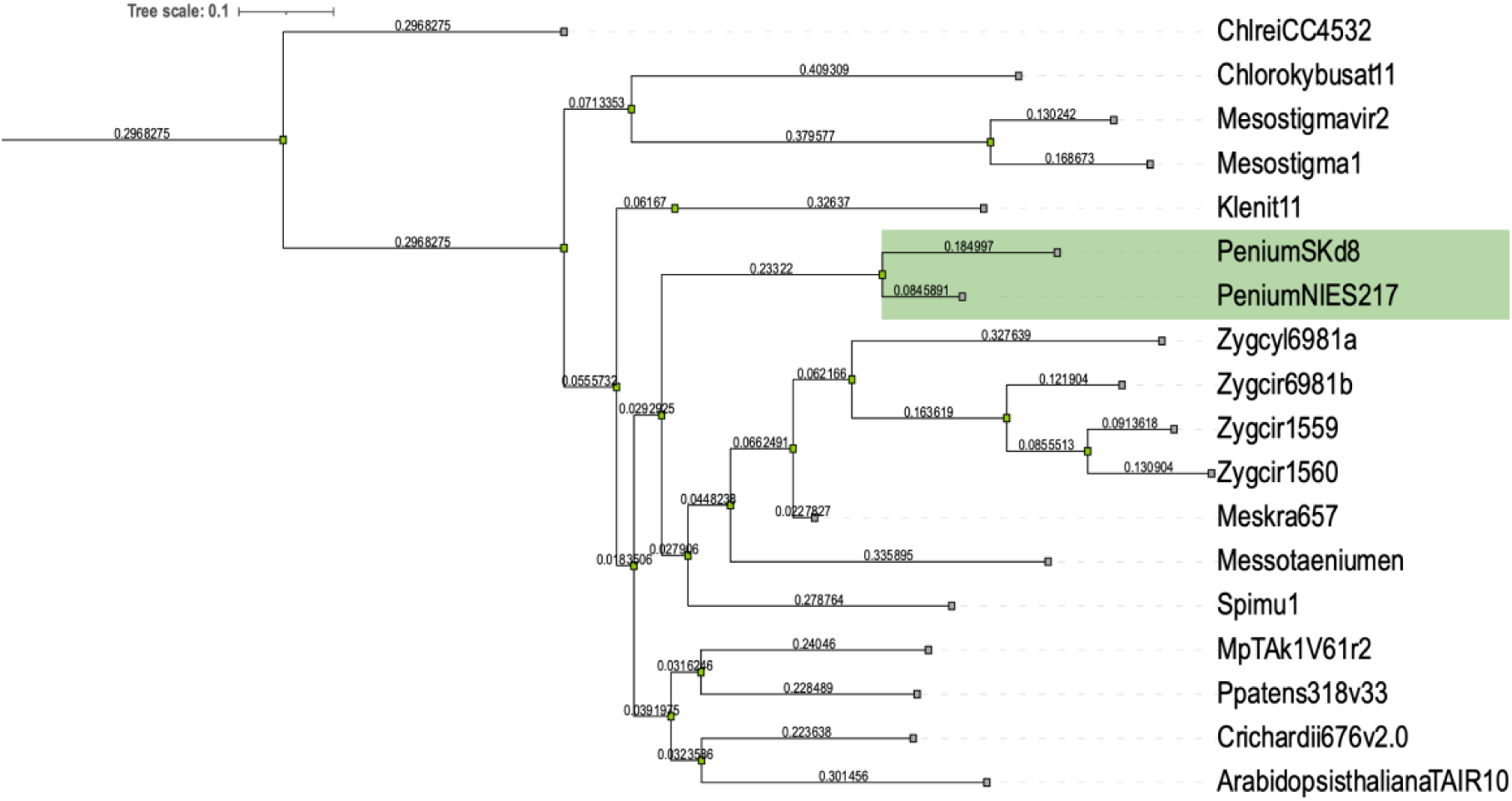
Species tree containing streptophyte algal species with available proteomes to place both P*enium margaritaceum* isolates PmSk#8 and PmNIES217. The tree was generated in Orthofinder using the complete proteome of all the contained species following the pipeline indicated on ^47^. Highlighted in green are both strains of *P. margaritaceum*.

**Fig S4.**
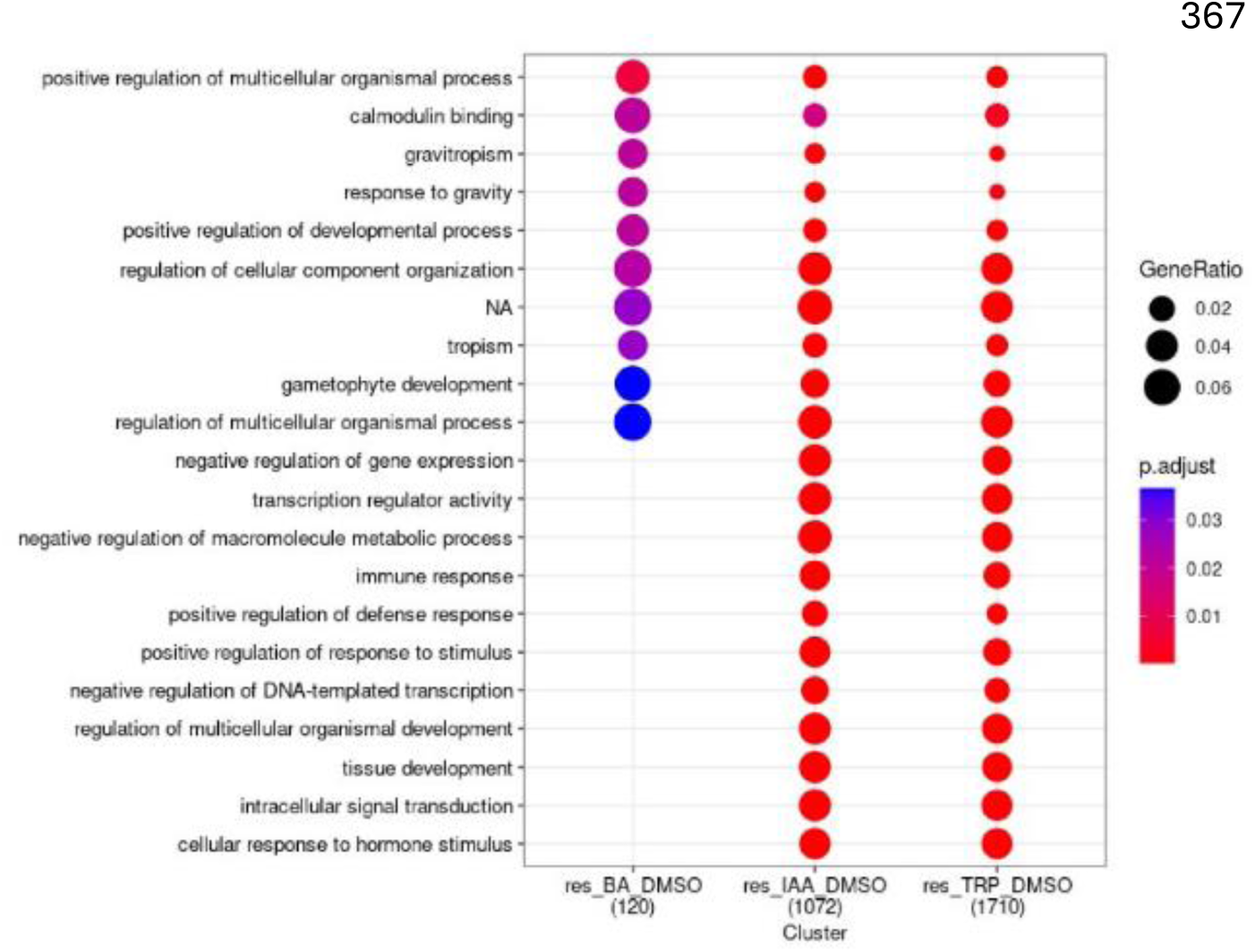
GO term analysis comparing functional enrichment between DEGs of Trp, IAA or BA-treated Penium cells with respect to DMSO.

**Table S1.** Summary of parameters used to quantify growth, division and changes in cytoplasmic streaming in Penium cells treated with different chemicals.

| Treatment | Average percentage of growing cells | Average percentage of dividing cells | Growth rate $\mu\text{m}/\text{h}$ (measured from intermittent growth) | Track mean speed $\mu\text{m}/\text{s}$ |
| --- | --- | --- | --- | --- |
| Mock | 18% | 6% | 0.025 | 18.6 |
| IAA | 41% | 31% | 0.033 | 22.5 |
| Trp | 66% | 45% | 0.035 | 25.8 |
| BA | 51% | 21% | 0.021 | 19.31 |

**Video S1. Growth of *Penium* cells in microfluidic devices over the course of 60 hours.** Annotations indicate that cells are in different growing stages. NG: non-growing, G:growing, D: dividing.

**Video S2. Penium cells stained with MitoTracker, treated with different chemicals showing cytoplasmic streaming.** Scale bar 10 µm.

## Methods

### EXPERIMENTAL MODEL AND SUBJECT DETAILS

#### Algal cell culture

*Penium margaritaceum NIES-217 (PmNIES217)* was obtained from the NIES Collection (https://mcc.nies.go.jp/), cells were grown in Woods Hole Medium (WHM) for 15 days ^48^, at 20 °C with long-day light conditions (16:8 hours), 30 – 50 µmol/m^2^/s in a shaker with continuous agitation (60 rpm) in 50ml or 250 ml Erlenmeyer flasks. Liquid cultures with an OD_750_ of 0.2-0.3 were used for experiments.

### METHOD DETAILS

#### Fabrication of devices for microfluidics

Microfluidic channels containing traps for *Penium* were fabricated using a mould from SU8-25 photoresist (Kayaku Advanced Materials) on a silicon wafer (Si-Mat, 76.2 mm diameter, P/Boron doped, orientation <100>). To prepare the mould, the wafer was first immersed in propylene glycol methyl ether acetate (PGMEA) and spin-coated for 30 seconds (s) at 500 rpm using a Laurell technologies spin coater until all PGMEA had evaporated. The wafer was then coated with SU8-25 photoresist and spin-coated for 30s at 500 rpm, followed by 90s at 2250 rpm, to obtain a 30 µm-thick layer. After coating, the wafer underwent a pre-expossure bake on a 65 °C hot plate O/N. to perform a pre-exposure bake, followed by exposure in a Microwriter ML3 Baby device (Durham Magneto Optics) at an exposure energy of 15 mJ/cm^2^ and a wavelength of 365 nm. Multiple copies of the design were arranged on one wafer. Post-exposure baking was carried out sequentially on a hot plate: 90 seconds at 65 °C, 270 seconds at 95 °C, followed by cooling for 2 hours at room temperature. The wafer was then developed by immersion it in PGMEA with gentle shaking for 5 minutes, followed by rinses with PGMEA, isopropanol and water. Finally, the mould was dried with compressed air and glued to a plastic Petri dish.

The devices were made from polydimethylsiloxane (PDMS) using a SYLGARD^®^184 silicone elastomer kit. The polymeric base and curing agent were mixed thoroughly at a 10:1 ratio, and centrifuged for 2 minutes at 50 x g to remove air bubbles. The mixture was poured onto the SU-8 mold, and the remaining air bubbles were removed with vacuum desiccation (15-40 minutes). The PDMS was then cured overnight at 70°C. After curing, the individual PDMS devices were cut from the wafer with a scalpel and peeled off. Holes for the tubing were made using a 1.5 mm diameter biopsy punch with plunger (KAI). Round glass cover slides with a thickness of 150 µm and diameter of 50 mm were prepared for bonding the PDMS chips by rinsing isopropanol, drying with compressed air and plasma cleaning for 1 minute. Both the cleaned glass slides and PDMS devices were plasma-activated for 15 seconds and immediately bonded by placing the surfaces together. The assembled devices were left to stabilize overnight at room temperature before use in experiments.

#### Auxin treatments in microfluidic devices

To prepare the microfluidic PDMS devices for experiments, the channels were flushed with 1% bovine serum albumin (BSA), filtered through a 0.22 μm membrane, and incubated for 5 minutes to coat the surfaces and prevent cell adhesion. The device was then mounted in a Nikon Ti2-E equipped with a Kinetix sCMOS camera. Diluted cultures, adjusted to a concentration of 1×102 cells/ml, were loaded in the PDMS traps using a continuous flow of WHM media chips at 300 bar pressure using an Elveflow® OB1 MK4 pressure controller coupled to an Elveflow® Mux Distributor distribution. Treatments in the devices were achieved by flushing WHM supplemented with DMSO as a mock control, 1 µM Indole-3-acetic acid (IAA), 0.1 µM IAA, 1 µM L-Tryptophan, 1 µM Benzoic acid. For growth experiments, WHM culture media was supplemented at a continuous flow for 60 hours using 50 mbar pressure. An automated tile experiment was set to record multiple tiles on different planes of the PDMS device. For cytoplasmic streaming measurements, cells were treated with corresponding chemicals for five minutes.

#### Imaging and treatments

For growth analysis, cells grown under continuous IAA, Trp, BA, DMSO treatments were imaged for 60 hours with 30 minutes intervals. Images were taken using transmitted light with a 10x objective in a Nikon Ti2-E equipped with a Kinetix sCMOS camera wide-field microscope.

For cytoplasmic streaming, treatments were achieved by treating cells for 5 minutes with IAA, Trp, BA, DMSO. Previous to imaging cells were stained overnight with MitoTracker Orange CMTMRos (Invitrogen™) to a final 500 nM concentration, protected from light, and agitated gently at 50 rpm. Excess dye was removed by washing the cells five times with WHM through centrifugation at 1620 x g before imaging and loaded into microfluidic chips. Movies of cytoplasmic streaming of cells were taken with a rate of 40 frames per second over 5 seconds using a 100X objective with oil immersion.

#### Cell growth image analysis

We used a cell segmentation algorithm optimized for *Penium* cells using MATLAB that automated the detection of cells and measurement of the length of cells in microscopy images, generating visual outputs to track the process and providing quantitative data to track cell growth and cell division (https://github.com/PoletC/PhD_thesis.git). The pipeline consisted of automated binarization of time series, followed by segmentation of identified objects (cells) using centroids. The length of the major axes of the centroid was used as a readout of cell growth. Finally, the pipeline includes a section for spatiotemporal cell tracking that links cell positions across time points. We plotted the relative growth points that correspond to the average length of the first five observations divided by the actual measurement of the length in an specific point. To distinguish the population of growing cells, we determine an experimental cut-off of 1.15 and a cut-off of 1.4 to determine the cells that divide.

Analysis of intermittent traces were performed according to the pipeline developed by ^28^. The interpolating growth traces were split into growing/non-growing regimes using a cut-off rate of 0.012 that was determined experimentally by fitting the growth curves of all the treatments (Figure S3). Only the growing cells were selected for individual observations. We tracked a total of 537 cells and performed four independent experiments per treatment.

#### Cytoplasmic streaming image analysis

Cells were stained overnight with MitoTracker Orange CMTMRos (Invitrogen™) to a final 500 nM concentration for cytoplasmic streaming measurements, protected from light and agitated gently at 50 rpm. Excess dye was removed by washing the cells five times with WHM through centrifugation at 1620 x g before imaging.

Recorded movies of cytoplasmic streaming were analyzed using the Fiji plugin Trackmate ^49,50^. Images were captured with a rate of 40 frames per second over 5 seconds using a 100X objective with oil immersion. Preprocessing was made using the “Subtract Background” tool, with a rolling Ball Radius of 80-100 pixels. We used the default Laplacian of Gaussian (LoG) detector for particle identification with an estimated object diameter of 1 micron and quality threshold adjusted depending on the image. The trajectories of the particles were generated using the default simple LAP tracker with a maximum linking distance of 5µm, two gap-closing maximum distances, and one frame-closing gap distance. The trajectories were then filtered to retain those with a duration of at least 3 seconds following the processing performed in *Arabidopsis* and *M. polymorpha* ^16^. Average particle speeds were computed and visualised in violin plots. Statistical differences between groups were performed using an ANOVA

#### RNAseq experimental setup and RNA extraction

Liquid cultures of *P. margaritaceum* were grown for 15 days under the conditions described above. After this initial growth phase, cells were collected by centrifugation at 1620 g for 5 minutes and re-inoculated in four 2 L Erlenmeyer for 6 days, with each flask representing a biological replicate. Large-scale cultures were maintained in an incubator (New Brunswick Innova 42) with controlled light and temperature settings to ensure uniform growth. Cells were harvested by centrifugation at 1620 x g for 5 minutes in 50 ml tubes. The cell pellets were washed five times with 10 mL of WHM medium to remove residual extracellular polysaccharides. The pellets were then resuspended in 1 ml of media and 200 µl of washed cells were re-inoculated in 10 ml of WHM supplemented with a final concentration of 1 µM IAA, 1 µM Trp, 1 µM BA or the same volume of DMSO. These cultures were incubated under the same controlled conditions for one hour.

Following the treatments, the cells were collected by centrifugation at 1620 x g for 5 minutes, immediately frozen in liquid nitrogen and freeze-dried to facilitate RNA extraction.

RNA was extracted using TRIzol™ (Thermo Fisher Scientific), with modifications to remove residual polysaccharides. Following the manufacturer’s recommendations 1 mL of TRIzol™ was added to the freeze-dried cell powder and incubated at room temperature for 5 minutes. To separate phases, 200 µL of chloroform was added, and the mixture was centrifuged at 12,000 × g for 15 minutes at 4°C. The clear supernatant was carefully transferred to a new tube. 500 µL of TRIzol™ was added to the supernatant, incubated for 5 minutes, and mixed with 100 µL of chloroform before centrifugation. This step was repeated to remove any remaining polysaccharides, which could affect RNA quality.

Then the upper phase was transferred to a new tube followed by the addition of 500ul of isopropanol for precipitation. The precipitate was purified using RNeasy spin columns (Qiagen) with for on column DNaseI digestion (Qiagen). 10 µl DNase I stock solution were added to 70 µl RDD buffer and added onto the column membrane. Digestion was performed for 20 minutes at room temperature. The column was then washed with 350 µl RWI-buffer, after discarding the flow-through the columns were washed with 500 µl RPE-buffer twice. RNA was then eluted with 40 µl of RNase-Free Water. For quality control, RNA samples were run in agarose gels, samples with intact bands were then run in a Bioanalyzer using an RNA 6000 Nano chip (Agilent Technologies) to determine RNA integrity and concentrations. RNA short-read (Illumina NovaSeq PE150) and long-read (PacBio Sequel II) library prep, quality control, and sequencing of were carried out by GenomeScan BV and Novogene Co., Ltd, respectively.

#### *De novo* annotation of the PmNIES 217 transcriptome

Raw reads were tested for quality control using FastQC. These raw reads were used to build the de novo assembly using Trinity with default parameters except "--SS_lib_type FR". Completeness of the new transcriptome was checked using BUSCO with ’viridiplantae_odb10’ database. Transcript sequences were converted to protein sequences using TransDecoder. Redundant protein sequences were removed using CD-HIT with a cut-off of 99% identity. Interproscan (v5.60-92.0) was used to annotate the protein domains, pathways and GO terms from the InterPro database. EggNOG mapper was used to map the gene ontology and orthogroup annotations from Viridiplantae (taxonomy ID: 33090) with percentage identity, minimum % of query and subject coverage all set to 30. Best-bidirectional BLAST hits from *Arabidopsis* (Araport11) and *M. polymorpha* (v6.1) were obtained by BLASTP to the proteomes. (location)

#### Differential expression and GO enrichment analysis

Raw read counts for the transcripts were obtained using Salmon (v1.9.0) through a pseudo-alignment approach based on the transcriptome generated and annotated earlier. Differential expression analysis was performed using DEseq2 package in R environment, with ‘∼ condition’ as the design matrix. Poorly expressed genes (with raw counts < 100) across all samples together were removed prior to the differential expression analysis. Principal components were calculated using the VSD normalized counts and plotted using ggplot2 package. Significant genes (padj<0.05 & log2(foldchange)>1) were obtained and further used for the downstream gene ontology (GO) analysis. GO enrichment was performed using the package ‘ClusterProfiler’ and dot plots were generated for the enriched terms (p<0.05). All the plots were generated using the ggplot2 package in the R environment.

### QUANTIFICATION AND STATISTICAL ANALYSIS

To evaluate the differences between treatment groups found in (Figures 2E, 4B), we performed a one-way ANOVA followed by Tukey HSD (Honestly Significant Difference) test (p<0.05) and followed by extracting the group labels. Statistics were generated using pandas and bioinfokit libraries in a Python environment.

